# Experimental sound exposure modifies swimming activities and increases food handling error in adult zebrafish

**DOI:** 10.1101/2021.12.01.470707

**Authors:** Reza Mohsenpour, Saeed Shafiei Sabet

**Affiliations:** University of Guilan; Behavioural Biology, University of Guilan

**Keywords:** Anthropogenic noise, Behaviour, Foraging performance, Sound impact, Zebrafish

## Abstract

Anthropogenic noise is increasing globally and is recognized as a source of environmental pollution in terrestrial and aquatic habitats. Sound is an important sensory stimulus for aquatic organisms and it may alter stress-related physiological indices and induce broad behavioural effects in a range of marine and freshwater fishes. Specifically, sound exposure may induce changes in swimming activities, feed efficiency and spatial distribution changes in fish. Here, we experimentally tested sound effects on swimming activities and foraging performance in thirty individually housed, captive adult Zebrafish (*Danio rerio*). Adult zebrafish and water fleas (*Daphnia magna*) were used as model predator and prey species, respectively. Acoustic stimuli consisted of four sound treatments with different temporal patterns. All had the same frequency range and were administered on average 121 dB re 1 µPa^2^/Hz. Our results constitute strong evidence for sound-related effects on zebrafish behaviour. All sound treatments led to a significant increase in the number of startle responses, and the brief and prolonged swimming speed for zebrafish. We found sound effects on the spatial distribution of zebrafish; Although there were no significant sound-related changes for horizontal spatial displacement in all treatments, zebrafish swam significantly more in the lower layer of the tank except during the irregular intermittent 1:1-7 in brief sound exposure treatment. The results of foraging performance showed that food discrimination error was unaffected by sound treatments and was low for the zebrafish. However, food handling error was affected by sound treatments; all treatments induced a significant rise in handling error. This study highlights the impact of sound on zebrafish swimming activities, and that more feeding bouts are needed to consume the same number of food items increasing energy demand under noisy conditions.

## 1. Introduction

Currently, due to increased human activities and global technology advancement since the Industrial Revolution, the Earth’s environment has undergone extensive changes (Normandeau Associates, 2012). These environmental changes can affect the planet itself and living organisms, which threatens global biodiversity (Kunc et al., 2016). The rapid rate of change poses many environmental challenges (Tuomainen and Candolin, 2011) in both terrestrial and aquatic habitats. For example, one of the main sources of environmental pollution, which may also can be recognized as an environmental stress stimulus, is anthropogenic noise that not only affects terrestrial animals, but also has many consequences on aquatic organisms (Popper et al., 2020; Slabbekoorn et al., 2010; Slabbekoorn and Ripmeester, 2008). Anthropogenic noise sources in aquatic habitats, such as military sonars, shipping activities, wind turbines, and pile-driving can occur frequently, having varying temporal patterns, and are widespread geographically (McDonald et al., 2006; Normandeau Associates, 2012). Consequently, anthropogenic noise has changed underwater soundscapes worldwide, represents a very subtle driver of environmental change and is a novel challenge to aquatic organisms.

Recent studies have investigated the impacts of anthropogenic noise on a wide range of taxa and across a variety of scales (Barber et al., 2009; Morley et al., 2014; Normandeau Associates, 2012; Slabbekoorn et al., 2010; Thomsen et al., 2021; Tyack, 2008). Anthropogenic noiseee can cause a range of physical, physiological, and behavioural disorders in aquatic organisms, including marine mammals (Erbe et al., 2018; Moore et al., 2012; Southall et al., 2008), seabirds (Bermúdez-Cuamatzin et al., 2018; Green et al., 2016; Hansen et al., 2020), reptiles (Injaian et al., 2020; Simmons and Narins, 2018), fish (Hastings and Popper, 2005; Hawkins, 1986; Mills et al., 2020; Popper et al., 2003), and invertebrates (Carroll et al., 2017; Coquereau et al., 2016; Murchy et al., 2019). Because of the high propagation rate and low attenuation over large distances in underwater environments, acoustic stimuli play an important role in aquatic environments (Tyack, 1998; Popper and Hastings, 2009; Slabbekoorn et al., 2010).

Behavioural effects are the most likely to occur and thus drive stress (Smith et al., 2004; Popper and Hawkins, 2019). Acoustic stimuli can act as a distracting stimulus (Popper and Carlson, 1998; Chan et al., 2010), interfere with detecting prey and antipredator behaviour (Hawkins and Myrberg, 1983), compromise foraging performance (Neo et al., 2015; Purser and Radford, 2011; Shafiei Sabet et al., 2015; Voellmy et al., 2016), disrupt reproductive behaviour (McCloskey et al., 2020), or mask important acoustic signals and cues for conspecific recognition and communication purposes (Codarin et al., 2009; Amorim et al., 2015; De Jong et al., 2018b; Hawkins and Picciulin, 2019) in a range of marine and freshwater species.

Many marine and freshwater fishes have well-developed hearing abilities that provide them with a key biological advantage to detect low intensities and perceive a broad range of frequencies (Hawkins, 1986; Heath et al., 2021; Popper et al., 2019; Wahlberg and Westerberg, 2005; Wysocki et al., 2006). While there are well-documented studies on how sound affects marine fish behaviour (de Jong et al., 2018a; Mortensen et al., 2021; Peng et al., 2015), and its adverse repercussions at the individual and community level, much less is known about how sound affects the behaviour of freshwater fish. To the best of our knowledge, few studies exist on this topic (Fedoroff, 2021; Mickle and Higgs, 2018; Pieniazek et al., 2020). Moreover, sound exposure can change spatial distribution and swimming behaviour of fish which may consequently affect foraging success if they avoid foraging in noisy food areas (de Vincenzi et al., 2021; Hanache et al., 2020; Hubert et al., 2021; Shafiei Sabet et al., 2016a). There is growing evidence that foraging performance is likely to be modulated by anthropogenic noise: instead of performing successful feeding bouts, fish can instead display typical behavioural stress responses (Purser and Radford, 2011; Shafiei Sabet et al., 2015; Voellmy et al., 2016). Currently, little is known about the effects of sound exposure on swimming activity and foraging performance of fish, although there are a few well-documented studies. It has been shown that increased boating activity was associated with a reduction in activity rates, changed vertical distribution and compromised foraging success of free-ranging mulloway (*Argyrosomus japonicus*) (Payne et al., 2015) and Mediterranean Damselfish (*Chromis chromis*) (Bracciali et al., 2012).

Other studies have shown that experimental sound exposure increases performance errors and therefore causes a negative impact on foraging efficiency in both the three-spined stickle back (*Gasterosteus aculeatus*) (Purser and Radford, 2011) and the European minnow, (*Phoxinus phoxinus*) (Voellmy et al., 2014a). More recently our previous study has also shown a clear sound impact on zebrafish foraging performance; more food handling errors occurred under noisy conditions (Shafiei Sabet et al., 2015). A primary consequence of sound exposure would appear to be a shift in the spatial displacement. The resulting disturbance might modify the allocated foraging time budget, foraging patterns, and the relative abundance of prey and predatory species. Such changes in turn may increase foraging energy demand and the amount of time fish must allocate to foraging which, subsequently may affect food searching, discrimination, and handling.

In general, *Danio rerio* is known as a model fish species in behavioural, environmental, and biomedical studies (Cachat et al., 2010; Egan et al., 2009; Whitfield, 2002). Zebrafish is a member of the Cypriniformes order and acclimates well in captivity (Detrich et al., 2011) and naturally lives in the tropical freshwater (Spence et al., 2008). Zebrafish have specialized hearing structures, the Weberian ossicles, with an auditory system that has homologies to the mammalian auditory system (Weber, 1820; Alexander, 1964; Whitfield, 2002).

Progress in the field of behavioural biology, conservation ecology, and toxicology is also associated with the study of invertebrates. Daphnia is a small crustacean that is an important part of the food web in freshwater habitats and inhabits many types of shallow water bodies (Ebert, 2005; Parejko and Dodson, 1991; Reynolds, 2011).

In the present study, which is a follow up to recent work (Shafiei Sabet et al., 20215), we experimentally tested the hypothesis that experimental sound exposure produced by an underwater speaker affects the general swimming activities and foraging behaviour of predatory zebrafish upon water flea prey under laboratory conditions. Compared to Shafiei Sabet er al., 2015 where we used an in-ear speaker, in this study we adopted an underwater speaker to increase sound pressure levels and induce a potential gradient in the arena. We also extended the time of behavioural observation to explore more long-term effects of sound on the zebrafish. Our specific goals were: first, to assess the effect of experimental sound exposure on zebrafish activities including swim speed and spatial displacement; secondly, to estimate whether the temporal pattern of sound exposure influences zebrafish behaviour; and third, to verify our recent laboratory-based findings of sound impacts on zebrafish swimming activity and foraging behaviour.

## 2. Materials and Methods

This study was performed in the ornamental fish breeding facility center at University of Guilan, Sowmeh Sara, Iran (37°17ʹ39ʺN, 49°19ʹ55ʺE), using a 50 ×15 ×20 cm aquarium between 10:00 and 14:00 every day. Thirty adult zebrafish (15 F: 15 M) were used and all zebrafish were approximately 45 days old and of the wild-type, short-fin variety. They weighed approximately (mean± s.d.) 1.23 ± 0.02 g) and were obtained from an ornamental fish breeding shop in Sowmeh Sara, Iran. Zebrafish were stored in a stock glass tank (50×30×40 cm) for two weeks and adapted to environmental conditions to reduce possible stress and hormonal changes from transportation and captivity conditions (Deakin et al., 2019). The fish were fed 0.8 mm commercial Biomar® feed until the day before the experiment (Neo et al., 2015). Water fleas were caught at the campus of the department in Sowmeh Sara every morning, during the entire experiment (See Shafiei Sabet et al., 2015; Shafiei Sabet et al., 2019).

### 2.1. Sound treatments

Four sound treatments with different temporal patterns along with a control treatment were used. The first treatment served as control treatment in which the fish were exposed to ambient noise (AN). Experimental treatments consisted of regular intermittent sound (IN) with a fast pulse rate (1:1) (Figure. 1 (a)), regular intermittent sound with slow pulse rate (1:4) (Figure. 1 (b)), irregular intermittent sound (1:1-7) (Figure. 1 (c)), and continuous sound (CS) (Figure. 1 (d)). All three intermittent treatments include one second of sound, but the difference between these sound treatments is the intervals between these sounds (silence time) (See also Shafiei Sabet et al., 2015). The sound treatments were generated and modified using Audacity® software (2.3.1) at the frequency range that zebrafish can detect(300-1500 Hz) (Higgs et al., 2002) as well as encompassing the bandwidth of anthropogenic sounds that overlap with the zebrafish’s hearing range, such as from vehicles, pump systems, and pile drivers (Slabbekoorn et al., 2010). Sound was produced by the software in the same frequency range (400-2000 Hz) based on settings from our earlier study (Shafiei Sabet et al., 2016).

**Figure 1:**
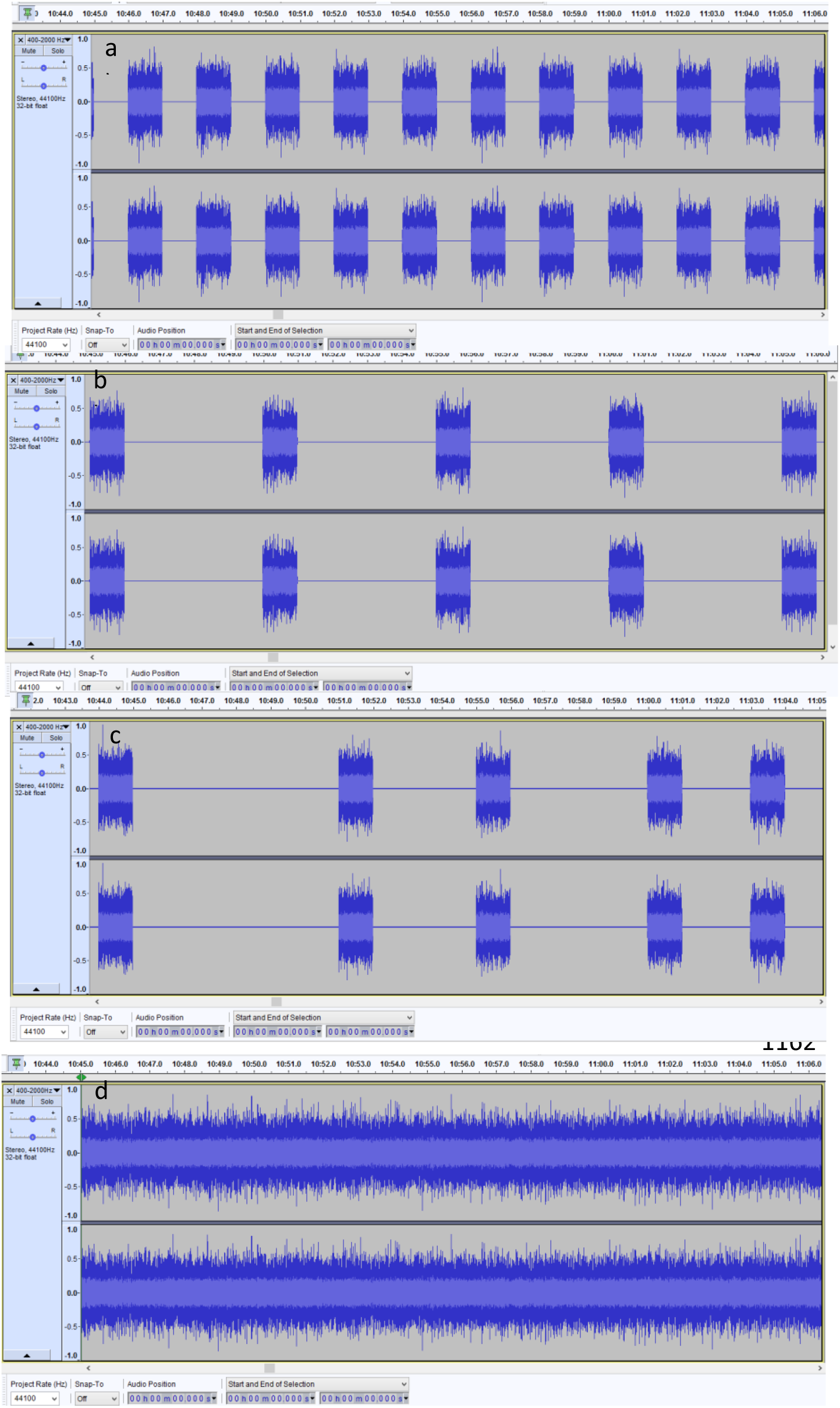
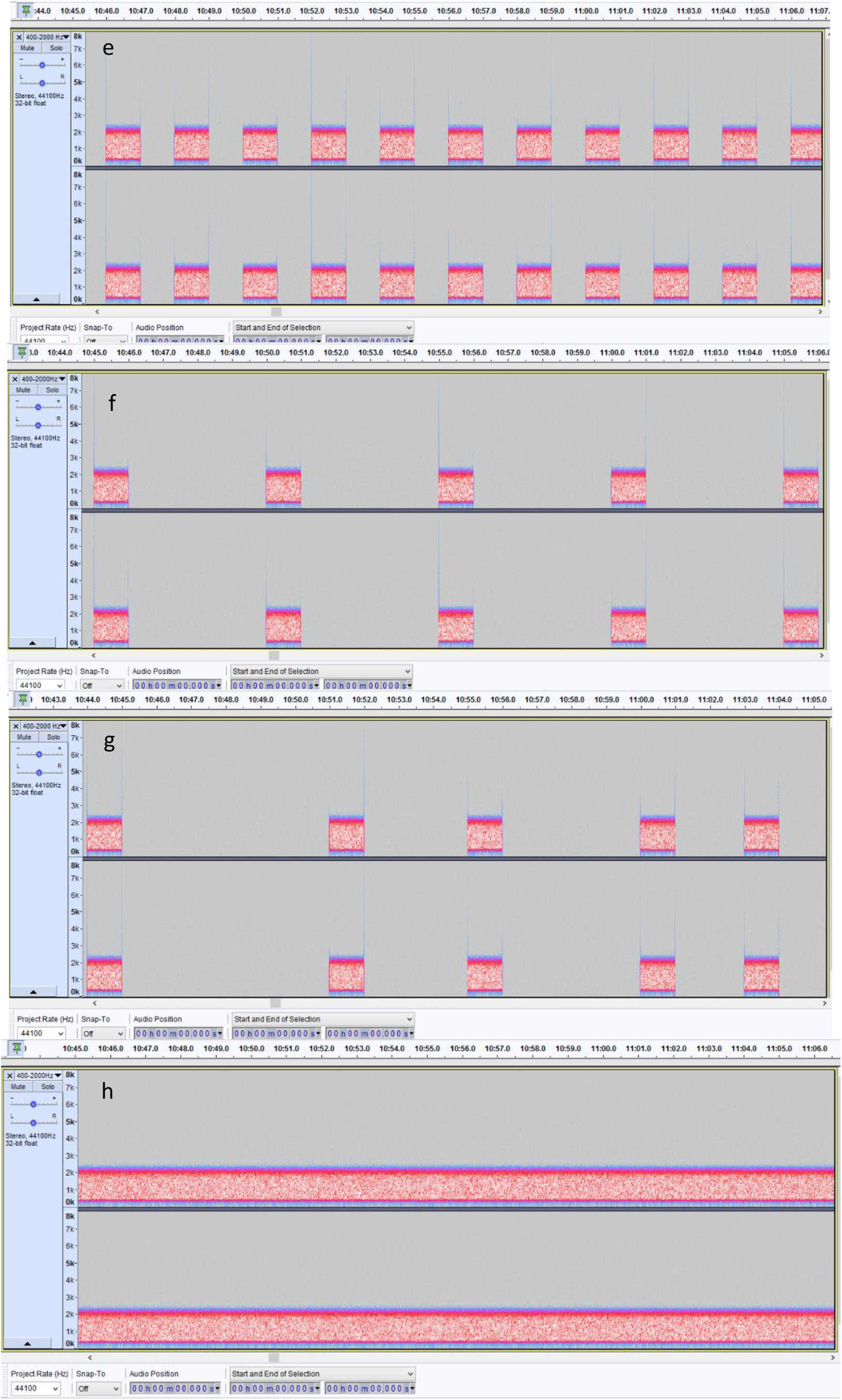
Waveforms and spectrograms (across a 20 s stime period) of all intermittent and continuous sound patterns used in the experiment (see also Shafiei Sabet et al., 2015). (a) Regular intermittent sound treatment with fast pulse rate (1:1), (b) Regular intermittent sound treatment with slow pulse rate (1: 4), (c) Irregular intermittent sound (1:1-7), (d) Continuous sound treatments accordingly.

### 2.2. Experiment Tank

The experimental tank (50×20×15 cm) was prepared with a black background to improve the contrast between Daphnia and fish in the video file. Zebrafish swimming activities were filmed by a video camera (Panasonic HC-V180 Full HD) at a distance of 50 cm from the front of the test tank. Sound treatments were played back as stereo WAV files using a Laptop (Sony Vaio SVF1421A4E) connected to an underwater speaker (custom-build speaker, 30 W, 10 Hz-10 KHz). In this experiment, a divider plate was placed in the tank to divide its length in half (25×20×15 cm) in order to restrict the zebrafish swimming environment and make the entire swimming space visible for video-recording (Figure. 2). The test tank was surrounded by black plastic so that the fish’s behaviour was not affected by other visual factors. Only the camcorder lens passed through the plastic, which was the same for all treatments and repetitions. To isolate the experiments from external, confounding sounds, we covered the walls at the entrance to the experimental room with egg boxes, both inside the room and in the hallway.

**Figure 2:**
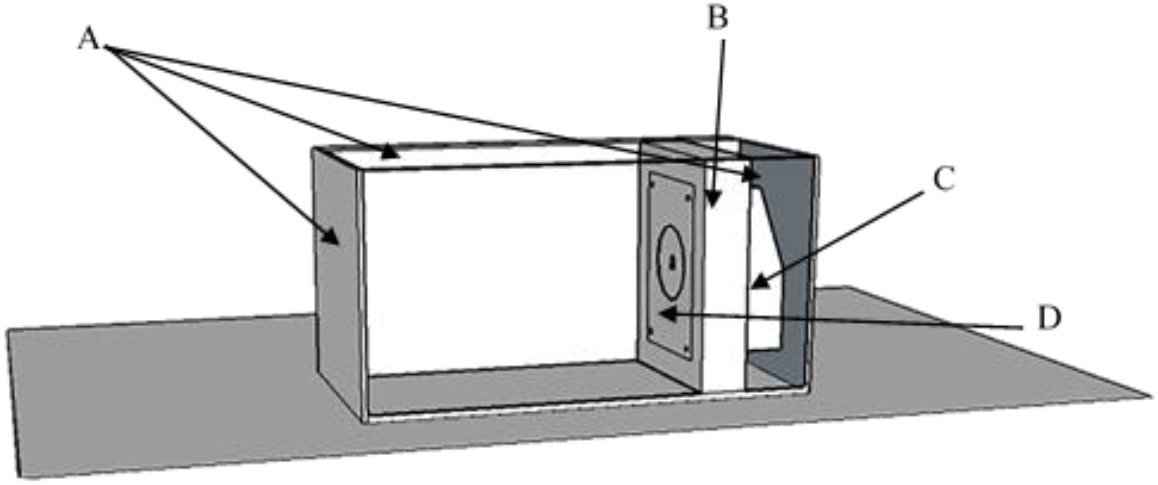
Experimental set-up. Schematic view of the test tank. A: Obscured pages of the test tank are used to enhance the fish’s visual contrast in the film. B: Underwater speaker space separated by screen from the fish swimming space. C: Underwater speaker holder box. D: Underwater speaker.

The underwater speaker used in the experiment was placed horizontally behind a separator plate (See Figure. 2). Then, each test was performed as follows: after ten minutes, the sound treatment was played by the speaker and sound player for 20 minutes. The food item (Daphnia) and a non-food item (Duckweed) were added to the experiment tank after ten minutes from the onset of each sound playback across all sound treatments. The fish were exposed to all five sound treatments on the same day, with a 15-minute interval between treatments. The procedure was repeated the next day with the next fish naïve to any sound exposure and no experience with sound treatments. The sound treatment playback order was randomized daily and balanced across days. This approach was previously described to quantify behavioural indices of zebrafish in response to acoustic measurements in this system (Shafiei Sabet et al., 2015). The photoperiod used in the experiment was 12:12 (Higgs et al., 2002; Villamizar et al., 2014) and the light intensity averaged 62 lux, measured by a light meter (TES_1336A – TES Electrical Electronic Corp. Taiwan). Water temperature and the amount of dissolved oxygen (DO) in the water were measured during the experiment and were maintained at 26±1 °C at 8±1 mg / L, respectively.

### 2.3. Underwater sound measurement in the tank

Sound files were played back using a sound player connected to a custom-build underwater speaker attached to a custom-build sound tuning amplifier (See Shafiei Sabet et al., 2015). The level of sound pressure was recorded by a hydrophone (model Aquarian Scientific AS-1) which was connected to the amplifier (model PA-4) and a Tascam linear PCM recorder (model DR-100MKII). Sound pressure levels and power spectral density of the sound for the continuous treatment were evaluated in R studio software (Version. 1.1.456). The sound pressure of continuous sound treatment during playback was on average 121 dB re 1 µPa^2^/Hz for 5 seconds and the ambient sound pressure was on average 96 dB re 1 µPa^2^/Hz for 5 seconds (Figure. 3).

**Figure 3:**
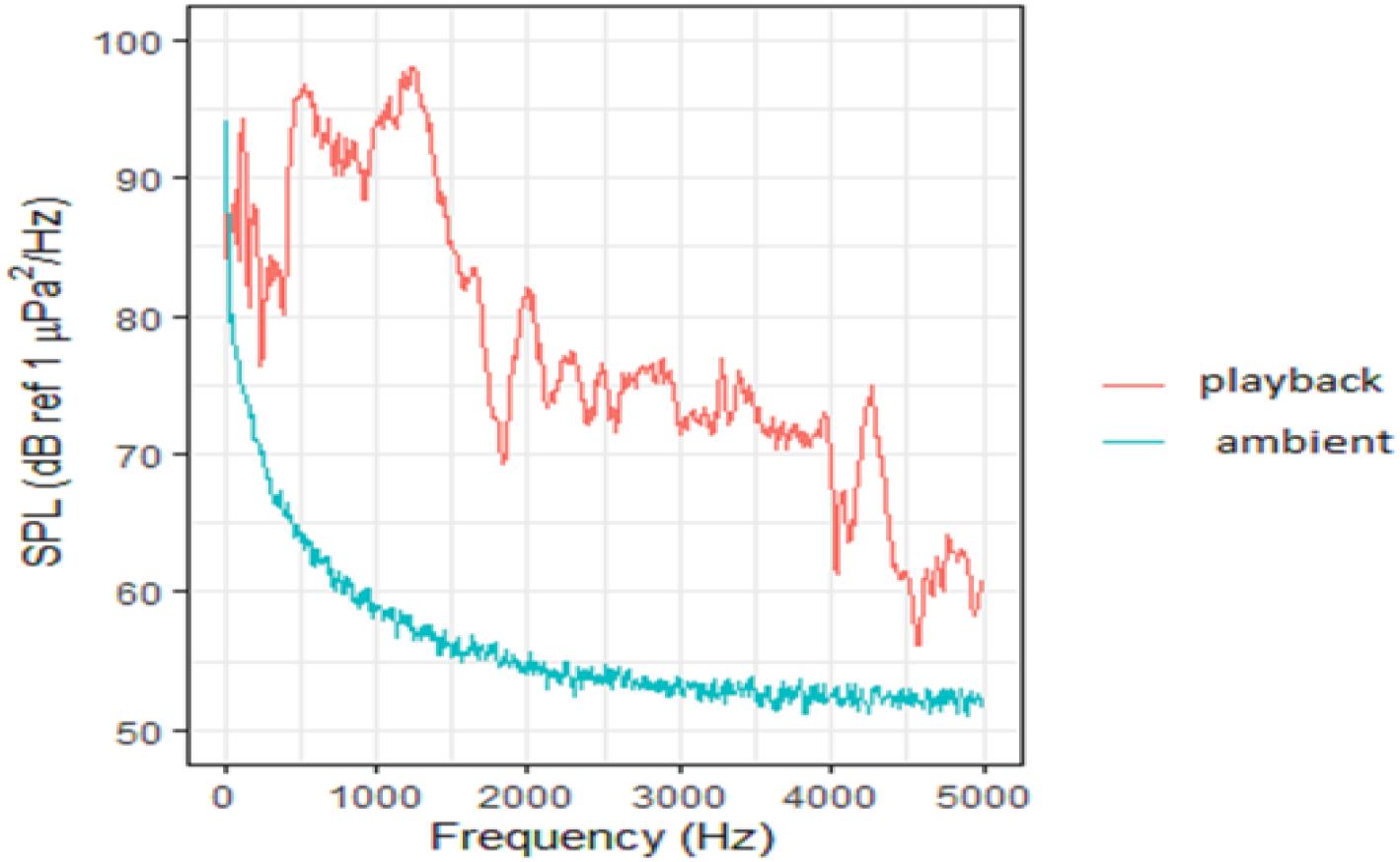
Spectral distribution of continuous sound pressure level compared to silent treatment (dB ref 1 µPa^2^/Hz). Silence conditions (blue) and continuous sound playback (red). The diagram shows that the sound intensity level has increased significantly in the range of hearing frequencies of zebrafish. For interpretation of the references to colour in this figure legend, the reader is referred to the web version of this article.

### 2.4. Effect of sound on Zebrafish swimming behaviour

To investigate the effect of sound on the behaviour of Zebrafish, all animals were introduced individually in the experimental tank. The fish were introduced to the test tank the day prior to testing (Overnighting) to acclimate to the environment so that they could clearly use the entire space of the tank for their swimming and display normal behaviour (Neo et al., 2015; Shafiei Sabet et al., 2015). All fish were given 30 minutes to acclimate in the experiment tank.

Zebrafish behavioural responses were video recorded for a maximum of 30 minutes for each treatment (10 mins before exposure and 20 mins during exposure). Swimming behaviour parameters such as the number of startle response (defined as the peak swimming speed at a rate of more than 10 cm per second that occurs immediately after the sound was played for one minute (See Shafiei Sabet et al. (2015)), brief swimming speed (during the last 5 seconds before sound exposure and 5 seconds with the onset of sound) and prolonged swimming speed (during one minute before the start of sound exposure and one minute with the onset of sound exposure) were evaluated for all treatments. Also, the spatial distribution of fish in the tank was explored. In the vertical dimension, tank height was divided into two parts: zero to 7.5 cm and 7.5 to 15 cm based on the 15cm dewatering height of the tank. To check the distance from the sound source in the horizontal dimension, the length of the tank was divided into three parts: zero to 8.33 cm, 8.33 to 16.66 cm and 16.66 to 25 cm.

### 2.5. Effect of sound on foraging behaviour of zebrafish on water fleas

The effect of sound on the foraging behaviour of zebrafish was investigated by mixing 5 water fleas (~3 mm; target prey species) and 5 non-food substances (~ 3mm non-food items; duckweed) were mixed in 25 ml beaker and adding the mixture gently to the fish tank. This procedure was done in the same manner for all treatments. The water fleas were in the same sizes that caught with plastic Pasteur pipettes with suitable entrance tube diameter to decrease damaging water fleas, which is suitable for feeding this species of fish at puberty and can be orally ingested by the fish (Shafiei Sabet et al., 2015). Both food and non-food items are naturally present in the habitat of this fish. Foraging performance was assessed as described in Shafiei Sabet et al., 2015.

### 2.6. Behavioural information processing and statistical analysis

Zebrafish behaviour videos were converted to 10 frames per second by Xilisoft Video Converter Ultimate software. This was done to reduce the magnification of time,, which increases the accuracy of the fish swimming survey by reducing the fish speed for spatial inspection per second. Logger Pro® video tracking software (Vernier Software, Beaverton, OR, USA, version 3.10.1) was used to examine behavioural responses including the number of startle responses, average swimming speeds, and spatial distribution changes of the fish. Data analysis was performed using SPSS 25 software and Excel 2016. The normality of the data was evaluated by Kolmogorov-Smirnov test and the homogeneity of the data was assessed using Levene’s test. Then, significant differences between the treatments were assessed with a repeated measures ANOVA analysis and using Tukey multi-range test. A Huynhe Feldt correction was performed when sphericity could not be confirmed in the repeated measures ANOVA. Bonferroni corrected post hoc tests were performed when ANOVA test results were significant. The level of significance in this study was considered P<0.05. A custom-written acoustic calibration script in R studio software (Version 1.1.456) was also used to evaluate sound pressure levels and power spectral density that were played by the underwater speaker.

## 3. Ethical note

All housing, handling and experimental conditions were in accordance with the guidelines for the treatment of animals in behavioural research and teaching (ASAB, 2020). Water fleas and zebrafish were allowed to acclimate gradually to laboratory conditions before they were used in any of the experiments and showed no signs of adverse effects from the experimental conditions. Zebrafish showed only a brief startle response upon onset of the moderate sound playbacks and no mortalities or physical damage were observed during experiments (Neo et al., 2015; Shafiei Sabet et al., 2015). There are no legal requirements for studies involving invertebrates such as waterfleas (Daphnia) in Iran.

## 4. Result

### 4.1. Impact of sound on swimming behaviour of Zebrafish

Experimental sound exposure changed zebrafish swimming activities in several ways. Sound treatments significantly increased the startle responses and swimming speed in fish (Figure. 4). The number of startle responses instantly increased upon exposure to sound so that all treatments showed a significant difference compared to the ambient condition (repeated measures ANOVA: *F*_3.23,93.68_=6.31, *P*<0.001). However, no significant differences were observed in terms of temporal patterns between sound treatments (*P*>0.05) (Figure 4a). Also, the brief swimming speed (5 seconds before the sound and 5 seconds during the sound) was significantly different in all treatments compared to the ambient condition (repeated measures ANOVA: *F*_3.06,88.70_=11.17, *P*<0.001), although there was no significant difference between the sound treatments with different temporal patterns for this measure (*P*>0.05) (Figure 4b). This difference in prolonged swimming speed (60 seconds before the sound and 60 seconds during the sound) was also impacted, so that there was a significant effect between sound treatments compared to ambient conditions (repeated measures ANOVA: *F*_3.39,98.34_=7.72, *P*<0.001). However, again there was no significant difference between the different sound treatments (*P*>0.05) (Figure 4C).

**Figure 4.**
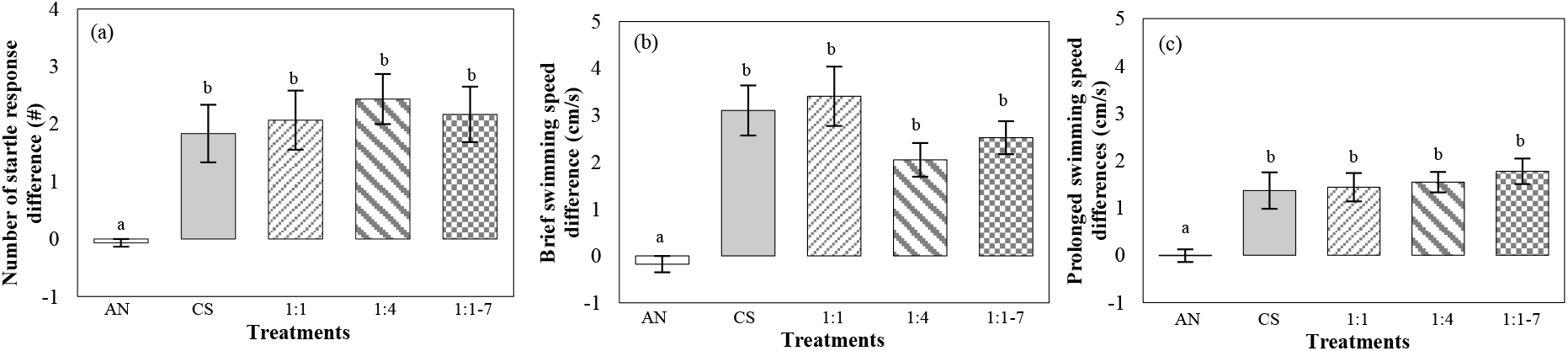
Effect of sound exposure treatment on swimming behaviour of Zebrafish. (a) Number of startle responses expressed as the difference between the first 60 seconds during sound exposure and the 60 seconds immediately preceding sound exposure: Fish were exposed to continuous sound (CS), three intermittent sounds (1:1, 1:4, 1:1-7), and ambient noises (AN) as control treatment (N=30, *F*=6.312, *P*<0.001, Standard error changes (±1)). (b) Brief swimming speed difference of Zebrafish between the first 5 seconds of sound exposure and the last 5 seconds preceding sound exposure for each sound treatment and the ambient condition (N=30, *F*=11.172, *P*<0.001, Standard error changes (±1)). (c) Prolonged swimming speed difference of Zebrafish between the 60 seconds preceding sound exposure and the first 60 seconds during sound exposure on all five treatments (N=30, F=7.725, *P*<0.001 Standard error changes (±1)).

Based on Figure 5, in all four sound treatments (CS, 1:1, 1:4, 1:1-7), a sudden increase in swimming speed was observed once sound treatments were played at the 60 second observation time. Observations also showed that in all sound treatments, fish swim speed returned to baseline after 60 seconds of playback. In ambient conditions (Figure 5a), no significant difference in swimming speed was observed during the entire time period. While in continuous sound and regular intermittent 1:1 treatment (Figure 5b+ 5c), a significant difference was observed when comparing the 5 seconds before sound playback and the first 5 seconds after sound playback (CS= repeated measure ANOVA: F_4.54,131.66_=9.53, *P*≤0.001, 1:1= repeated measure ANOVA: F_3.85,111.72_=11.72, *P*≤0.001). However, for both treatments, the fish returned to their baseline speed during the last 5 seconds of sound exposure: there was no significant difference in swim speed between the 5 seconds preceding sound exposure and the final 5 seconds of playback (*P*>0.05). This pattern was also observed in the regular intermittent 1:4 and irregular intermittent 1:1-7 treatments (Figure 5d+ 5e) with a more intense difference before and during the sound (1:4= repeated measure ANOVA: F_6.24,181.01_=14.17, *P*≤0.001, 1:1-7= repeated measure ANOVA: F_7.14,207.15_=15.01, *P*≤0.001).

**Figure 5.**
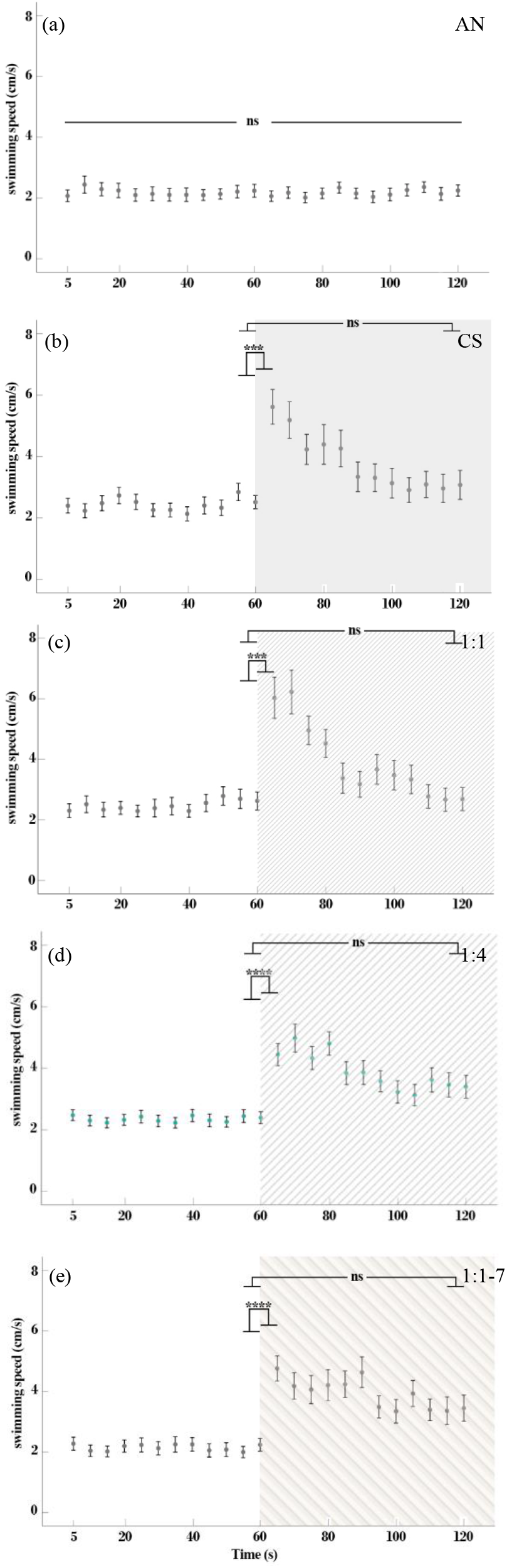
Effect of sound treatments on zebrafish prolonged swimming speed: (a) Ambient condition, (b) continuous sound, (c) regular intermittent 1:1, (d) regular intermittent 1:4, (e) irregular intermittent 1:1-7. The time was divided into three period bins for formal statistical analysis: the last 5 seconds before sound exposure, the first 5 seconds during sound exposure and the last 5 seconds during sound exposure. Prolonged swimming speed decreased with repeated exposure in all sound treatments. (NS= no significance, ***= *P*≤0.001, ****= *P*≤0.0001).

Like the continuous and 1:1 treatments, no significant difference was observed between the swim speed of the 5 seconds preceding sound exposure and the last 5 seconds of the treatment: the fish returned to their baseline speed (*P*>0.05).

Changes in the spatial distribution of zebrafish were investigated when exposed to sound treatments in vertical scale (lower layer) (Figure. 6) and horizontal scale (X position) (Figure. 7). According to Figure 6a, there was no effect on the average percent of time spent in the lower layer of the tank during sound treatments exposure over a brief time (15 seconds before sound and 15 seconds during sound exposure) (*F*_3.77,109.47_=1.486, *P*=0.214) or nor was there an interaction of treatment*times (*F*_3.55,102.92_=0.634, *P*>0.05). However, there was a main effect of times (*F*_1,29_=28.274, *P*<0.001). In the ambient (AN) and irregular intermittent sound (1:1-7) treatments, there was not a significant effect of time (AN= *F*_1,29_=28.274, *P*>0.05, 1:1-7= *F*_1,29_=28.274, *P*>0.05). However, there was a significant difference in two sound treatments before and during sound exposure: continuous sound (CS) and intermittent 1:4 (CS= *F*_1,29_=28.274, *P*<0.001, 1:4= *F*_1,29_=28.274, *P*<0.001). Also, in the intermittent 1:1 treatment there was a significant difference before and after sound exposure (*F*_1,29_=28.274, *P*<0.01). According to Figure 6b, the average percent of time spent in the lower layer of the tank during sound treatments over a prolonged time (60 seconds before sound and 60 seconds during sound exposure), was not affected by treatment (*F*_3.65,105.92_=1.837, *P*>0.05) or the interaction of treatment*times (*F*_4,116_=1.780, *P*=0.137). However, there was a significant effect of times (*F*_1,29_=35.398, *P*<0.001). In control treatment (AN) there was not a significant difference before and after sound exposure (*F*_1,29_=35.398, *P*>0.05). While in the continuous treatment (CS) and intermittent 1:4 treatment, there was a significant difference before and after sound exposure (CS= *F*_1,29_=35.398, *P*<0.001, 1:4= *F*_1,29_=35.398, *P*<0.001). The intermittent 1:1 treatment was also significantly different before and after sound exposure (*F*_1,29_=35.398, *P*<0.001). Also, the irregular intermittent sound treatment (1:1-7) showed a significant difference before and after sound exposure (*F*_1,29_=35.398, *P*<0.05).In contrast, there was no significant difference in the horizontal profile (X position) between sound treatments and control conditions, both in the brief and prolonged time (*F*_4,116_=1.369, *P*>0.05) (Figure 7a), (*F*_3.78,109.71_=1.810, *P*>0.05) (Figure 7b). Further, according to Figure 8 the effects of different sound patterns on the spatial distribution of fish can be seen as a heat map in brief (figure 8a) and prolonged duration (figure 8b). Acoustic treatments affected changes in swimming pattern from the top layer to the bottom layer but had no effect on the distance from the sound source located on the right side of the tank.

**Figure 6.**
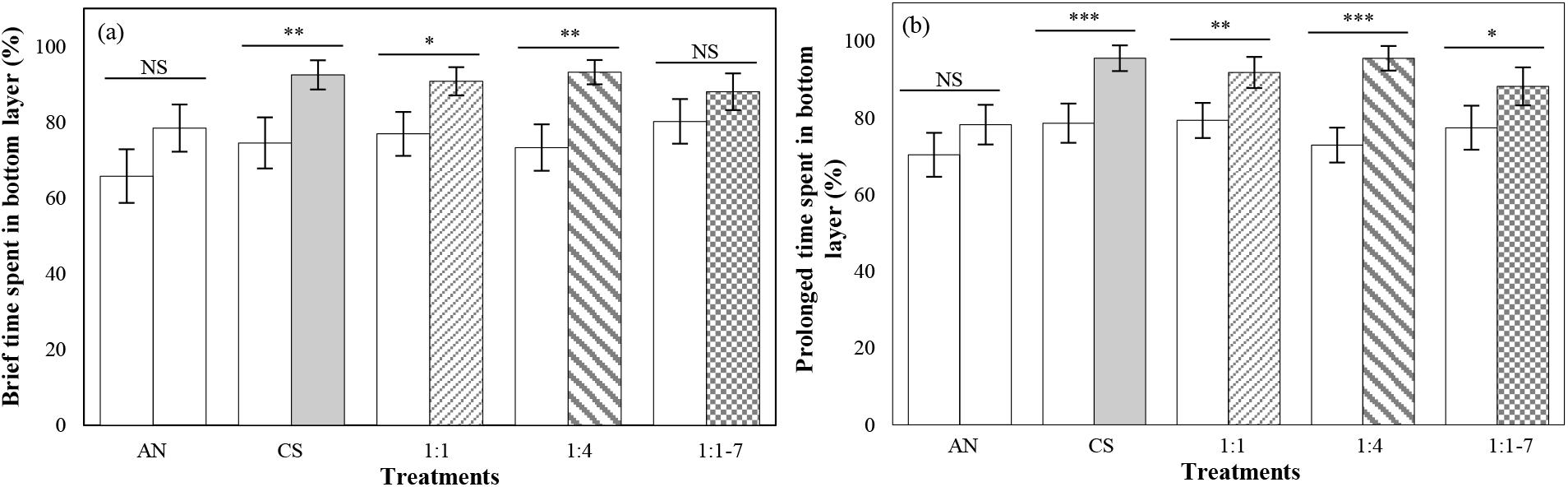
Average percentage of fish time spent in bottom layer of tank (N=30). (a) Brief time (15 seconds before and 15 seconds after sound exposure) (NS= no significance, *= *P*≤0.05, **= *P*≤0.01). (b) Prolonged time (60 seconds before and 60 seconds after sound exposure) (NS= no significance, *= *P*≤0.05, **= *P*≤0.01, ***= *P*≤0.001). Bottom layer area for spatial displacement was defined as the bottom layer up to 10 cm from the bottom of the tank. (df= 1) Standard error changes (±1).

**Figure 7.**
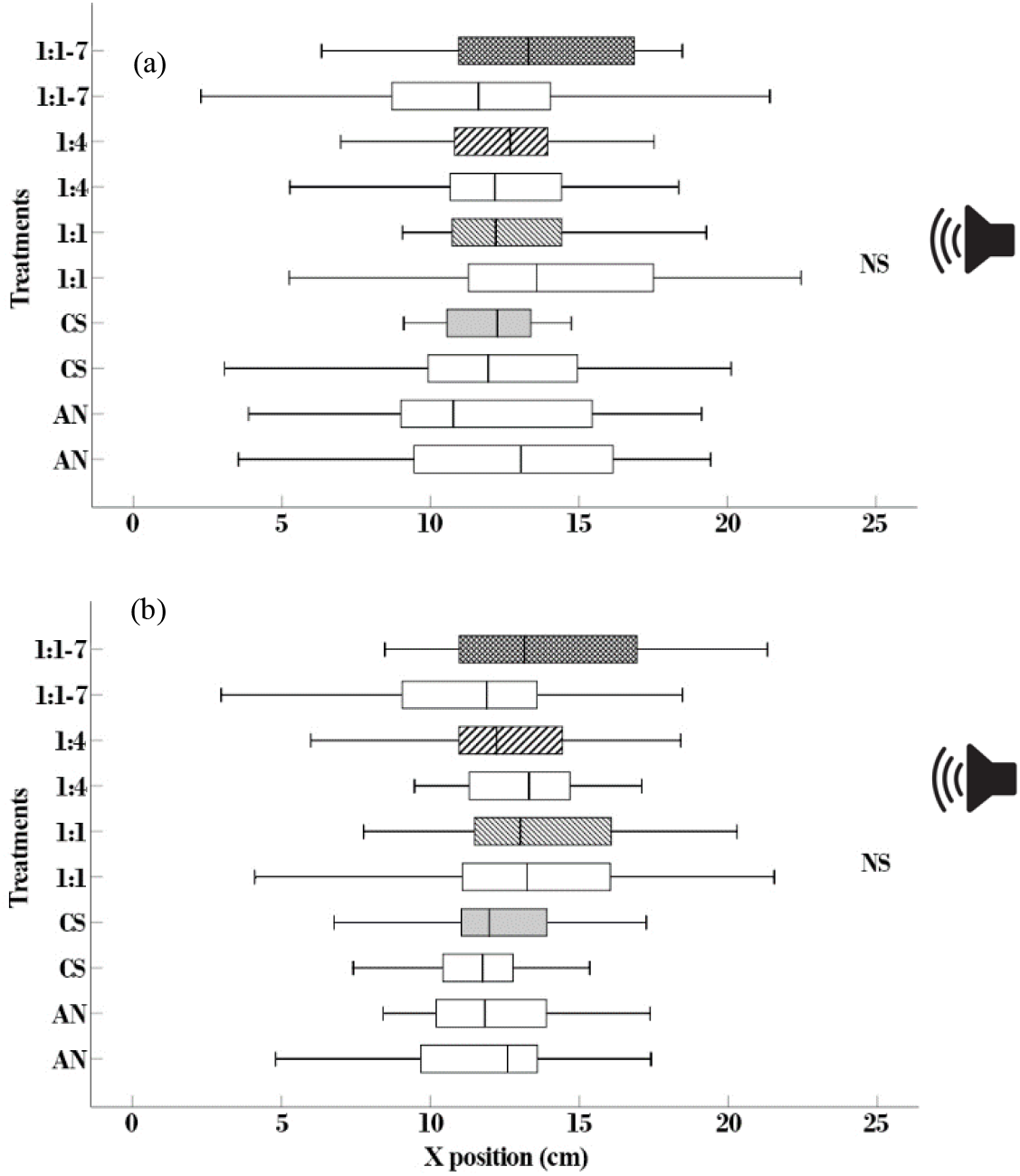
Effect of sound exposure on horizontal spatial distribution of Zebrafish. (a) Brief time (15 seconds before and 15 seconds after sound exposure). (b) Prolonged time (60 seconds before and 60 seconds after sound exposure). The underwater speaker played back from the right side of the tank. Bars show Means ± SE.

**Figure 8.**
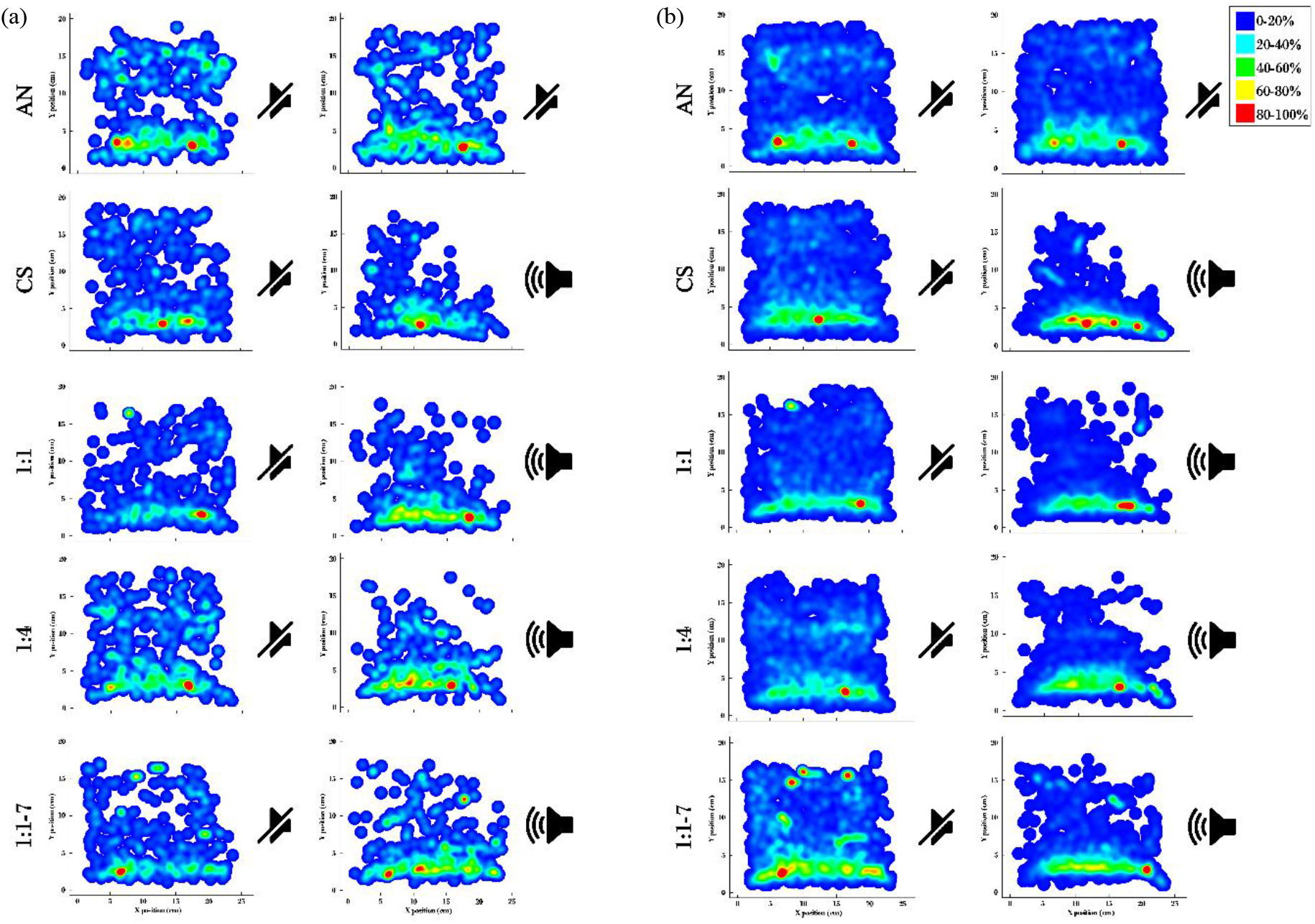
Heat map of fish swimming in the tank environment. (a) Brief time (15 seconds before and 15 seconds after sound exposure). (b) Prolonged time (60 seconds before and 60 seconds after sound exposure). The blue color (#0000FE) indicates the 0-20 % of the fish in the tank. The aqua color (#01FFFF) indicates the 20-40 % of the fish in the tank. The lime color (#00FF01) indicates the 40-60 % of the fish in the tank. The yellow color (#FFFF01) indicates the 60-80 % of the fish in the tank. The red color (#FE0000) indicates the 80-100 % of the fish in the tank. The underwater speaker played back from the right tank. For interpretation of the references to colour in this figure legend, the reader is referred to the web version of this article.

### 4.2. Impact of sound on foraging performance of Zebrafish

According to Figure 9a, none of the acoustic treatments had a significant difference compared to the ambient noise treatment on zebrafish food discrimination error (repeated measure ANOVA: F_4,116_=1.339, P>0.05). In fact, there was no food discrimination error between food and non-food item by broadcasting sound treatments compared to control treatment.

**Figure 9.**
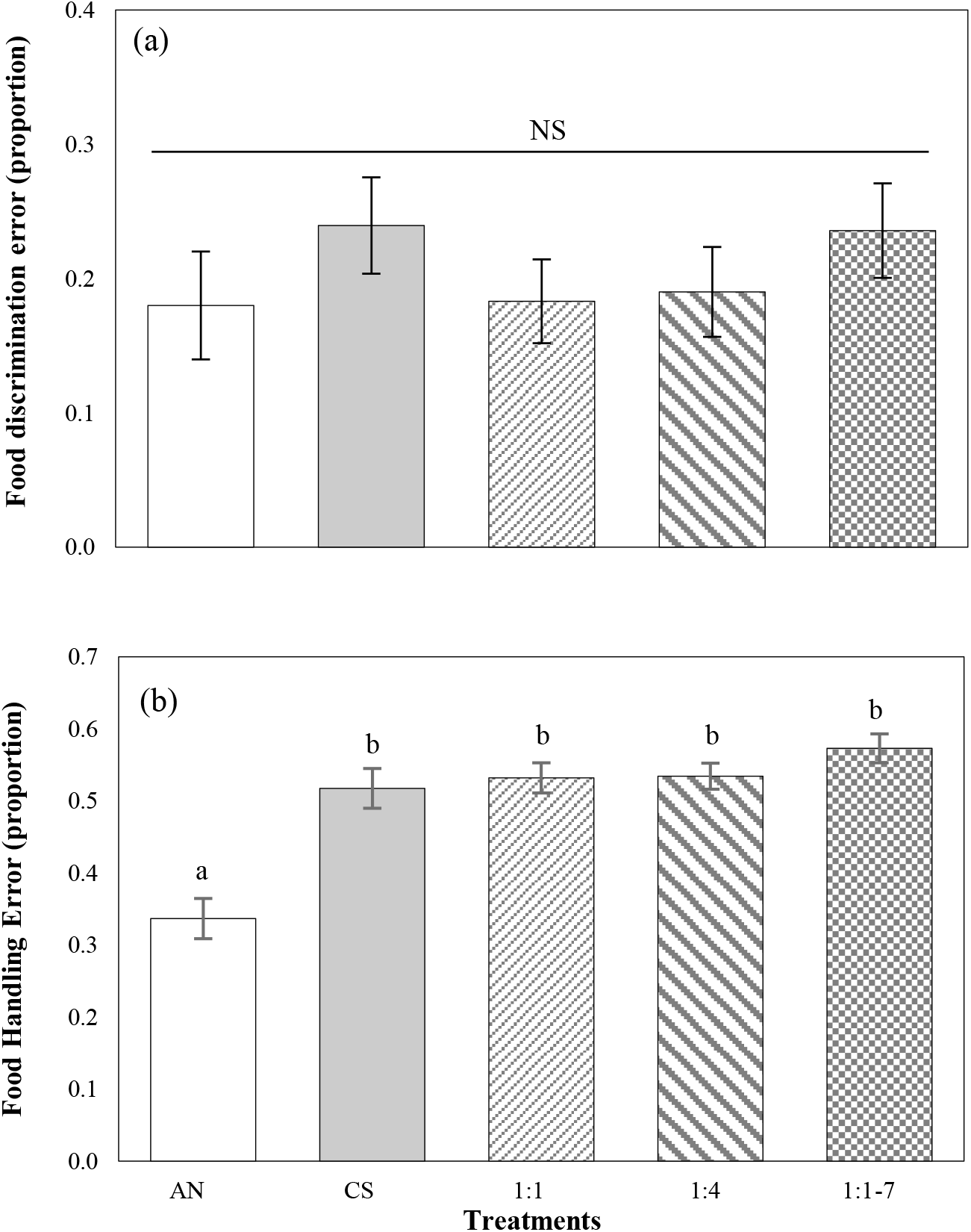
Effect of acoustic stimulus on foraging activities of Zebrafish. (a) Food discrimination error; it is described as the proportion of duckweed particles attacked relative to the total number of attacks to both duckweed particles and water fleas from the introduction of food items until the end of sound exposure in sequence for each zebrafish individual (Shafiei Sabet et al., 2015). (N=30, df=3.756, *F*=1.339, *P*=0.226) (b) Food handling error; It is described as the proportion of the total of water fleas attacked that were missed or released again after initial grasping from onset of food introduction until the end of sound exposure in sequence for each zebrafish individual (Shafiei Sabet et al., 2015). (N=30, df=2.825, *F*=26.023, *P*=0.000019). Standard error changes (±1)

However, for food handling error, all acoustic treatments showed a significant difference compared to the control treatment (repeated measure ANOVA: F_2.82,81.91_=26.023, P≤0.001). There was no significant difference observed between sound treatments though (P>0.05) (Figure. 9b).

## 5. Discussion

Our results unequivocally demonstrate that acoustic stimuli affect zebrafish behaviour in different ways. In this experiment, zebrafish swimming behaviour indices such as the number of startle responses, the difference in brief swimming speed (within 5 seconds), the difference in prolonged swimming speed (within 60 seconds), and the spatial distribution in response to continuous and intermittent sound patterns were examined. The number of startle responses, indicative of anxiety in zebrafish and other aquatic species (Blaser et al., 2010; Maximino et al., 2010), in different sound treatments were significantly different compared to the control condition. However, these changes did not show a significant difference between sound treatments. Also, there was a significant increase in in brief and prolonged swimming speeds in all sound treatments compared to the control condition. The increased swim speeds were not significantly different between sound treatments though. Moreover, in support of previous work (Shafiei Sabet et al., 2015), this data showed the same impact of sound exposure on foraging performance in zebrafish. All will be discussed further in the following sections.

### 5.1. Startle responses as a specific indicator of moderate anxiety in zebrafish

The startle response is an involuntary action that is controlled by a pair of brain neurons in the Mauthner (M-) cells in the mesencephalon and plays a major role in the decision-making process (Eaton et al., 1977; Eaton et al., 1991; Mirjany et al., 2011; Zottoli, 1977). Increasing the intensity of the sound triggers an involuntary and very sudden C-startle response by Mauthner cells in the mesencephalon, leading to an involuntary escape response in fish (Eaton et al., 1977; Eaton et al., 1991). The fish’s increased number of startle response and brief swimming speed in response to sound treatments, are due to behavioural responses related to fear and anxiety in this species. Previous studies have shown that sounds increase motor acceleration and startle responses in zebrafish (Neo et al., 2015; Shafiei Sabet et al., 2015). However, Shafiei Sabet et al. (2015), using an in-air speaker as a sound source, reported no significant differences in the startle response of fish who were exposed to continuous or intermittent (1:1) sound compared to ambient noise, which is not consistent with the results of this study. The reason for this discrepancy in stress-related swimming behaviour could be due to differences in ambient noise intensity before the test, differences in the method of ensonifiying the fish tank (under water compared to in-air speaker), different housing conditions for the fish, and genetic differences and individual characteristics of the zebrafish.

Other species of fish respond similarly in response to sound exposure. European minnow (*<u>Phoxinus phoxinus</u>*) and sticklebacks (*<u>Gasterosteus aculeatus</u>*) showed a significant increase in the number of startle responses when exposed to sound, similar to the results of this study (Purser and Radford, 2011; Voellmy et al., 2014a). A startle response at the onset of sudden sound exposure is a common behavioural reaction in fish kept in captivity and in laboratory conditions. Of course, fish in open water and natural conditions can also show behavioural responses related to fear and anxiety (Neo et al., 2016; Staaterman et al., 2020). Spiga et al. (2017) stated that European seabass (*Dicentrarchus labrax*) also showed a higher number of startle responses under continuous and intermittent sound treatments compared to an ambient treatment.

A startle response by prey fish is a behavioural adaptation to enhance survival rate in the context of predator-prey relationships (Webb, 1986). When hearing the sounds of predator fish and those related to an attack, the prey fish will explosively swim in the opposite direction of the perceived sound in order to increase their chance of escaping from the predatory species. Sound can affect the prey fish’s decision-making power against sources of danger, and how they assesses risk (Dukas, 2004). However, it can also act as a distraction and cause inappropriate responses to danger (Chan et al., 2010; Simpson et al., 2015). It has been suggested that increased sound levels can potentially impair the perception of danger by predatory fish species (Slabbekoorn et al., 2010). It has been suggested that involuntary and acquired behavioural responses related to fear and anxiety are associated with the potential for the presence of danger (Blaser et al., 2010; Maximino et al., 2010). The quality and quantity of behavioural responses of captive fish to brief and severe stress stimuli are different from those of fish living in a naturalistic habitat: the behavioural responses can be amplified in captivity compared to what is seen in the wild (Malavasi et al., 2004). One reason for these differences could be due to the ability of most wild fish to respond before the signal reaches the stimulus threshold, so that wild fish have a longer time to make decisions and escape from potential danger. They also have more space available than in controlled laboratory environments. Another reason could be the high level of basal stress potential in controlled laboratory environments, which paired with additional stress from perception of the predator species can lead to more intense responses.

In addition to behavioural responses, increasing sound levels can also affect physiological responses in the laboratory and in the natural habitat of fish. The study of Spiga et al. (2017) and Radford et al. (2016) showed that sound exposure had a significant effect on the number of opening and closing of gills and thus on the gill ventilation of European seabass (*Dicentrarchus labrax*) compared to the control treatment. This increase in oxygen demand by European bass (*Dicentrarchus labrax*), which is accompanied by an increase in gill ventilation and the opening and closing of gill operculum, indicates an increase in stress levels. Santulli et al. (1999) showed that blood biochemical parameters including cortisol and glucose in European seabass (*Dicentrarchus labrax*) increased in sound treatments compared to the ambient treatment. Staaterman et al. (2020) stated that anthropogenic noise treatments in the natural environment also have the potential to affect stress-related physiology in coral reef fish, so that the amount of cortisol in sound treatments was significantly increased compared to the control treatment.

### 5.2. Sound impacts on the behavioural tolerance in swimming activities

In past work performed on zebrafish at the group and individual level, different sound patterns had a significant effect on zebrafish swimming activities such as the brief and prolonged swimming speed of the fish compared to the ambient treatment, which is consistent with the observations in the present study (Neo et al., 2015; Shafiei Sabet et al., 2015; Shafiei Sabet et al., 2016a). In the experiment of Neo et al. (2015), groups of 5 zebrafish had increased mean swimming speed during the intermittent (1:1) treatment compared to ambient and other sound treatments. Similar results were also previously observed in cod fish (*Gadus morhua*) and European seabass (*Dicentrarchus labrax*), which was consistent with the results of this study (Handegard et al., 2003) (Neo et al., 2018). The prolonged swimming speed of zebrafish weakened with repeated sound exposure suggesting a habituation process. Zebrafish might experience negative consequences if they are disturbed and do not habituate accordingly to sound exposure. Shafiei Sabet et al. (2016b) compared the effect of sound exposure on the swimming behaviour of two species of The Lake Victoria cichlids (*Haplochromis piceatus*) and zebrafish (*Danio rerio*). That study showed that the application of sound treatments reduced the swimming speed of cichlids (*Haplochromis piceatus*) and increased the swimming speed of zebrafish (Shafiei Sabet et al., 2016b). The reason of the difference in the swimming speed of cichlids (*Haplochromis piceatus*) can be related to species-specific behavioural traits in response to acoustic stimuli, genetic characteristics, and habitat conditions.

### 5.3. The effect of sound on the spatial distribution of zebrafish

Studies on the stress indices of zebrafish in the face of different sound patterns showed that with the onset of sound treatments, the spatial distribution of fish changes and the fish shows a greater tendency to swim in the lower layer of the aquarium environment. Neo et al. (2015) revealed that zebrafish showed startle response and increased brief swimming speed upon the start of sound treatments, and their spatial distribution changed so that fish was more inclined to swim in the top layers of the test tank. Also, there were no observations of freezing or standing in the lower layer of the tank (Neo et al. 2015). However, the results of the present experiment showed that zebrafish spent more time in the lower layer of the test tank, which is contrary to the report of Neo et al. (2015). One of the reasons for this vertical distribution difference could be the amount of sound intensity emitted by the treatments used in these two studies. The intensity of sound previously emitted in acoustic treatments was equal to 112 dB re 1µ Pa, which is less than the intensity of sound in this study (121 dB re 1µ Pa). Therefore, potentially a variety of increase in sound intensity between treatments and ambient noise can lead to different responses in fish.

It is also important to consider the potential impacts of acoustic stimuli in real field conditions. Other fishes have also shown spatial distribution changes in response to acoustic stimuli in field conditions. Neo et al. (2018) found that anthropogenic sounds increase the swimming depth of European seabass and distance from the sound source, which is consistent with the present study in terms of vertical distribution results. In a field study (Kok et al., 2021) have shown that bottom-moored echosounders, representative of a high intensity impulsive intermittent anthropogenic noise, affect the abundance, schooling cohesion behaviour, and swimming depth of pelagic fish. Two recent telemetry tagging studies demonstrated the effects of another intermittent source of sound, seismic surveys, on free-ranging benthic fish species. Bruce et al. (2018) showed that anthropogenic noise cause changes in diurnal local activity patterns and general swimming speed shifts of marine fishes and therefore may negatively affect the catchability of marine fish catchability and the. van der Knaap et al. (2021) demonstrated in a field study that Atlantic cod (*Gadus morhua)* left noisy areas more quickly and decrease their swimming activity compared to the ambient sound treatment. They also have shown that seismic surveys can disrupt diurnal activity cycles which may have long term negative consequences on this species’ foraging success and fitness. Although to date, there is a gap in knowledge in terms of freshwater soundscape environments and how elevated sound levels affect fish in freshwater habitats (Bolgan et al., 2016; Linke et al., 2018; Rountree et al., 2020; Putland and Mensinger, 2020).

Swimming towards the upper layer of the tank at the beginning of the sound treatment has been considered to be driven by curiosity and searching behaviour. This is based on past work where low-pitched sounds were emitted during routine feeding by care staff, which attracted the fish’s attention and caused them to gather at the top of the tank in anticipation of a meal (See Shafiei Sabet et al., 2015), However, the opposite response seen here, where fish gathered at the bottom layer of the tank during sound treatments, indicates the occurrence of stress and fear in fish, which is similarly expressed in studies on other fish species (Neo et al., 2018; Sarà et al., 2007). Responses to other stimuli including chemicals and fear extract have shown that fish move to the lower layer of the tank upon exposure and this spatial distribution response is interpreted as a behavioural indicator of fear in many fish species (Gerlai et al., 2000; Gerlai et al., 2006). Another reason for the contrasting results observed in Neo et al. (2015) and the present study could be the difference in ensonifying underwater using speakers: an in-air speaker was used in the previous study (Neo et al., 2015) and an underwater speaker was used in the present study. The use of in-air speakers to broadcast sound treatments leads to more slightly intense sound in the deeper parts of the tank than in the middle and the surface layers, which may cause the fish to move and escape towards the surface layer where less sound intensity is felt (Shafiei Sabet et al., 2015). However this observed spatial behaviour change with the onset of sound exposure may be also an exploratory behaviour and curiosity of fish (c.f. Neo et al., 2015)

The spatial distribution data found in this study is not consistent with what was found previously (Shafiei Sabet et al. (2015), despite using treatments with similar sound intensity (122 dB re 1µ Pa). The reason for this difference in spatial distribution could be the use of in air speaker in the study by Shafiei Sabet et al. (2015) and underwater speaker in this study, as well as the complexity of sound distribution patterns and sound gradients in aquarium environments (Campbell et al., 2019). Other factors such as fish tank size and available arena, fish life stage, and location of the fish storage tank, as well as differences in the species, genetics, and personality of the fish could contribute to the differing results. However, no significant differences were observed in the horizontal distribution by both Shafiei Sabet et al. (2015) and the current study.

Based on available sources for measuring sound intensity and scattering patterns in aquarium environments, Parvulescu (1967) and Akamatsu et al. (2002) demonstrate the complexity and variability of sound scattering patterns and sound gradients in enclosed aquarium tank environments. This indicates a major limitation of studying the spatial distribution of aquatic animals in enclosed and controlled environments that must be considered. Therefore, in order to study the distribution patterns of fish and other aquatic species more accurately, it is recommended to conduct field studies in the species’ natural environments in order to obtain a more comprehensive understanding of sound-dependent distribution patterns in aquatic species.

### 5.4. The importance of particle motion in fish tanks; behavioural observations for future works

In order to understand the behavioural changes of zebrafish in response to acoustic stimuli, first of all, it is very important to understand the mechanisms behind and that how the species detects and processes, and how it behaviourally responds to sound (Hawkins and Popper, 2020). Because well-documented studies already showed that the auditory system of fishes evolved primarily to detect particle motion, many fishes are most sensitive to particle motion and they can use it to determine the direction of a sound source (Hawkins and Popper, 2018; Popper and Hawkins, 2018; Sand and Bleckmann, 2008; Sisneros and Rogers, 2016). Underwater sound consists of both sound pressure and particle motion (Popper and Hawkins, 2018). In fact, all fishes and all invertebrates primarily detect particle motion but not all detect sound pressure. Only some of fishes, including the zebrafish, are sensitive to sound pressure as well as the particle motion (Popper and Fay, 2011; Popper and Hawkins, 2018). There are some studies revealing directional hearing and sound source localization in fish under laboratory conditions and in free sound fields. Schuijf (1975) proposed that the cod (*Gadus morhua*) are able to detect sound directionality by monitoring the particle motion of the sound field, presumably employing the directional orientation of the inner ear sensory cells (Dale, 1976). Although, Schuijf (1975) also concluded that to determine the direction of a sound source, the direction of only particle motion may not be sufficient. It has already been shown that cod could discriminate between signals coming towards the head as compared to those coming towards the tail (Buwalda et al., 1983; Schuijf and Buwalda, 1975). They argued that to eliminate any remaining 180^0^ ambiguities, directional hearing might involve both comparing the responses of hear cells oriented in different directions and also analysis of the phase relationship between the sound pressure and particle motion components (Schuijf, 1976).

### 5.5. How important particle motion is to fishes and invertebrates

We did not mention the levels and direction of the particle motion that is generated within the fish tank because of our restriction in acquiring the particle motion measurement devices due to the COVID-19 pandemic. Therefore, we believe it is premature to conclude that zebrafish cannot localize sound source in our experimental set up. One reason might be because we know very little about hearing in only over 120 of the more than 33000 known fish species (Ladich and Fay, 2013) and that the empirical and theoretical work on sound source localization and directional hearing in fishes have been contradictory and obscure for decades (Sisneros and Rogers, 2016). Further reasoning can be attributed to practicality: it is difficult to monitor particle motion in a fish tank, there is a lack of easily used and reasonably priced instrumentation to measure particle motion, there is a lack of sound exposure criteria for particle motion, and finally there is a lack of particle motion measurement standards (Popper et al., 2014).

Within an aquarium tank the levels of particle motion are often highest at the water surface, and close to the tank walls, when an underwater loudspeaker is used (Jones et al., 2019). However, resonant frequencies and reverberation may influence propagation and spectro-temporal structure of the received acoustic stimuli in fish ranks (Jones et al., 2019). Our fish moved towards the lower levels of the tank, which may be because the particle motion levels were highest close to the water surface, and lower at the bottom of the tank. It is always important to monitor the particle motion when examining the effects of sounds upon fishes and invertebrates (Nedelec et al., 2016; Popper and Hawkins, 2018). Moreover, invertebrates are especially sensitive to substrate vibration (Aimon et al., 2021; Hawkins et al., 2021; Morley et al., 2014; Roberts et al., 2016), and some fish are too. Particle motion measurement may play an important role in answering crucial biological and ecological questions relating to fishes and other species among taxa (Nedelec et al., 2016). Thus, in doing future experiments to explore anthropogenic noise impacts on the behaviour of fishes or invertebrates under laboratory conditions it is necessary to develop open source and accessible protocols for monitor both particle motion on three axes and sound pressure.

### 5.6. Acoustic stimuli trigger negative effects in zebrafish foraging performance

In the present study, the parameters of fish foraging behaviour such as food discrimination error and food handling error were examined. The results showed that the zebrafish did not show any significant difference in food discrimination error when exposed to sound treatments compared to ambient noise. Also, the food handling error data showed that all sound treatments were significantly different compared to the ambient treatment, but these sound treatments were not different from each other. Here we confirmed our earlier findings (Shafiei Sabet et al., 2015) that sound impacts go beyond a single species. Experimental sound exposure caused more food handling errors and foraging in zebrafish, which led to more waterflea survival in noisy conditions.

Previously, sound treatments significantly affected the foraging performance of sticklebacks: food discrimination error and food handling error increased significantly and foraging performance reduced significantly compared to the ambient treatment (Purser and Radford (2011). In terms of food discrimination error, this is not consistent with the results of the present experiment. One reason for this difference could be due to physiological differences between the fish species and their diets. Particularly, potential differences in the fish’s visual abilities could be one of the factors influencing these differences. In terms of food handling error, a similar result was previously observed with those of the present experiment. In other studies, by Voellmy et al. (2014b), there was a significant difference in the number of unsuccessful takes of Daphnia in stickleback and no significant difference in minnow fish, which can generally indicate the physiological difference between the two species and possibly the difference in visual sense between the two species. The minnow fish, which belongs to the Cyprinidae family and is similar to zebrafish family, did not show a significant difference in the unsuccessful takes of Daphnia.

In a study almost similar to the present experiment, Shafiei Sabet et al. (2015) designed an experiment to investigate the effect of different sound patterns on the swimming and foraging behaviour of zebrafish and found that the application of sound treatments caused a significant difference in fish handling error. There was no significant difference in food discrimination error, which was consistent with the results of this experiment. Sound playback can influence the ability to identify a food substance, and make decisions about attacking it. Since zebrafish have a strong visual sense and use this modality for hunting, sounds can affect the ability to perceive potential vision potentially via attentional shifts and distraction. Field studies have also focused on the effects of sound on other fish species. In a comparative study between locations in the Laurentian Great Lakes and in laboratory conditions, Pieniazek et al., 2020 showed that experimental white noise and boat noise changed swimming patterns and foraging behaviour of freshwater fish species in different ways and with highly variable patterns according to family. They demonstrated that captive black bullhead, *Ameiurus melas*, foraged less and startled more when exposed to both white noise and boat noise treatments. Furthermore, escape behaviour has also been demonstrated in elasmobranches in response to anthropogenic noise (Mickle et al., 2020; Mickle et al., 2022). In a laboratory-based study on two fish species, the spot tail shiner (*Notropis hudsonius*) and the bluegill sunfish (*Lepomis macrochirus*), with different hearing abilities it has been shown that shipping noise caused similar but also specie specific responses (Stasso et al., 2023). Our laboratory experiments were conducted to explore usefulness of laboratory based experiments to assess anthropogenic noise impacts and address fundamental behavioural research questions. We hypothesised questions to verify that acoustic stimuli affect swimming behaviour and foraging performance of captive zebrafish as a model species and establish insights and new hypotheses for future exploration to be conducted on other freshwater fish species in captivity and in the field.

## 6. Conclusion

The results of this study highlighted impacts of acoustic stimuli on a freshwater fish species and confirmed our earlier study findings on the same fish species (zebrafish) under laboratory conditions. Our findings show that zebrafish swimming parameters and foraging behaviour in various sound treatments were significantly different compared to ambient noise conditions. Sound pollution as a stressor in the brief and prolonged time can cause behavioural changes and disturbances in individual aquatic species and have broad and important repercussions on the communities of an ecosystem. It should be noted that the results of this study are obtained in captivity and under laboratory conditions therefore the interpretation of the results should be done with caution and attention should be given to the conditions of natural environments and behavioural limitations of any species.

## Funding

The current research was funded by the Iranian Ministry of Science, Research and Technology, University of Guilan with the grant number: 971.511.210004.

## Appendices

## Data accessibility

All data used for the analyses reported in this article and some videos are available from the figshare. Moreover, the data and some videos that support the findings of this article are available from the first author upon request.

## Declaration of competing interest

The authors declare that they have no known competing financial interests or personal relationships that could have appeared to influence the work reported in this paper.

## Acknowledgments

Thanks to Jeroen Hubert and Ozkan Sertlek for their guide and advice on hydrophone calibrations, editing R script versions and acoustic measurements. Special tanks to Hans Slabbekoorn for providing accessibility to acoustic measurement devices. We also thank Narjes Karimi for helping us in reviewing the results and graphs.

## Notes

### Competing Interest Statement

The authors have declared no competing interest.

### Summary of Updates

Updated discussion part

